# Baltic *Methanosarcina* and *Clostridium* compete for electrons from metallic iron

**DOI:** 10.1101/530386

**Authors:** Paola Andrea Palacios Jaramillo, Oona Snoeyenbos-West, Carolin Regina Löscher, Bo Thamdrup, Amelia-Elena Rotaru

**Author notes:** Present address: Department of Microbiology and Molecular Genetics, Michigan State University, Michigan East Lansing, United States (Oona Snoeyenbos-West).

## Abstract

Microbial induced corrosion of steel structures, used for transport or storage of fuels, chemical weapons or waste radionuclides, is an environmental and economic threat. In non-sulfidic environments, the exact role of methanogens in steel corrosion is poorly understood. From the non-sulfidic, methanogenic sediments of the Baltic Sea corrosive communities were enriched using exclusively Fe^0^ as electron donor and CO_2_ as electron acceptor. Methane and acetate production were persistent for three years of successive transfers. *Methanosarcina* and *Clostridium* were attached to the Fe^0^, and dominated metagenome libraries. Since prior reports indicated *Methanosarcina* were merely commensals, consuming the acetate produced by acetogens, we investigated whether these methanogens were capable of Fe^0^ corrosion without bacterial partners (inhibited by an antibiotic cocktail). Unassisted, methanogens corroded Fe^0^ to Fe^2+^ at similar rates to the mixed community. Surprisingly, in the absence of competitive bacteria, Baltic-*Methanosarcina* produced six times more methane than they did in the mixed community. This signifies that Baltic-*Methanosarcina* achieved better corrosion alone, exclusive of an operative bacterial partner. Our results also show that together with acetogens, *Methanosarcina* interact competitively to retrieve electrons from Fe^0^ rather than as commensals as previously assumed.

## INTRODUCTION

The Golf of Bothnia of the Baltic Sea has been the dumping ground for chemical weapons and radionuclide waste sheltered by steel containers [1–3]. Thus, corrosion of metallic iron (Fe^0^) structures is a health and environmental threat. Microbial Induced Corrosion (MIC) accounts for 20% of total corrosion costs just when considering calamities and prevention for the oil and gas industries alone [4,5]. MIC has been largely studied in marine environments (sulfide-rich) where sulfate-reducing bacteria cause rapid corrosion [6]. However, in non-sulfidic environments like the Bothnian Bay, corrosion of infrastructure may occur due to cooperative interactions between microorganisms such as methanogens and acetogens [7–11]. Microbial associations are critical to our understanding of MIC because cooperating partners apparently adapt and interact with Fe^0^ simultaneously, promoting corrosion rates above those they would induce as single species [12]. However the role of cooperative interactions in corrosion is understudied. Methanogens in general, and *Methanosarcina* in particular, were suggested to play an important role in corrosion, and were found associated with corroded structures from oil, gas, sewage water storage and transportation facilities [7–11], but also in aquifers where radionuclide-waste is stored underground [13].

Nevertheless, very few highly corrosive methanogens have been described (*Methanococcus maripaludis* strains KA1, Mic1c10 and MM1264 and *Methanobacterium* strain IM1) yet none belongs to the genus *Methanosarcina*. Studies on corrosive *Methanococcus* and *Methanobacterium* methanogens indicated that their high corrosive potential could not be explained by the small amount of H_2_ generated abiotically on Fe^0^ [14–18]. Consequently, the mechanisms proposed for these different methanogenic strains employed either 1) a direct uptake route [17,19] or 2) an extracellular enzyme-mediated electron uptake [20,21].

First, direct electron uptake from Fe^0^ or electrodes has been suggested as an alternative to abiotic-H_2_ uptake for *Methanobacterium* strain IM1 because:

1. It generated more methane from Fe^0^ than a H_2_-utilizing *Methanococcus maripaludis* strain [19] with a low H_2_-uptake threshold [22]
2. It produced methane using only a cathode poised at – 400mV under conditions unfavorable for abiotic H_2_ evolution [17].

However, we do not know in what way IM1 reclaims electrons directly from Fe^0^ or electrodes, or if other methanogens have similar abilities.

By inference we propose that *Methanosarcina* might be able to retrieve electrons directly from Fe^0^. This is because we recently reported that *Methanosarcina barkeri* could retrieve electrons directly from a poised cathode at −400 mV under conditions unfavorable for abiotic H_2_-evolution [23]. The same *Methanosarcina* was previously shown to grow on Fe^0^ (although assumedly using H_2_) [24] retrieve electrons from electrogenic syntrophic partners [25,26] or via electrically conductive particles [27–29]. During the interaction of *Methanosarcina* with an electrogenic syntroph, the two are in an obligate metabolic cooperation with one another [26]. As such, only *Geobacter* is provided with an electron donor - ethanol, and the methanogen with an electron acceptor – CO_2_. *Geobacter* is a respiratory bacterium that demands an electron acceptor to oxidize ethanol [30,31]. The electron acceptor may be extracellular (e.g. electrodes or cells [25]) then *Geobacter* uses their extracellular electron transfer machinery (outer membrane c-type cytochromes and electrically conductive pili [26,32]). During their interaction with *Methanosarcina, Geobacter* uses the cell-surface of the methanogen as electron acceptor. The methanogen is also favored by the interactions because it can utilize the electrons to reduce CO_2_ to methane. Only recently, plausible scenarios for direct electron uptake in *Methanosarcina* were substantiated using a comparative transcriptomic approach [33]. That study compared the transcriptome of *Methanosarcina* provided i) directly with electrons from an electrogen (*Geobacter*) or ii) with H_2_ from a fermentative-*Pelobacter*. Several redox active cell-surface proteins were specifically up-regulated in the *Methanosarcina* grown via direct electron uptake but not via H_2_-uptake [34]. However, the exact role of these cell-surface proteins in direct electron uptake by *Methanosarcina* is unidentified, and remains to be characterized.

The second described strategy for methanogens to reclaim electrons from Fe^0^ is by using extracellular enzymes in order to effectively capture electrons [18,35]. For effective electron recuperations, enzymes like hydrogenases, formate dehydrogenases or the heterodisulfide reductase supercomplex use Fe^0^-derived electrons to produce H_2_ or formate [18,35]. If an extracellular enzyme-dependent strategy would be useful in environmental-corrosive communities is yet to be determined. This is especially relevant, because, when sacrificial populations discharge extracellular enzymes they lead to enzymatic-H_2_/formate to be taken advantage off by unspecific and diverse H_2_/formate-utilizers. Moreover, outside the cell the stability of sensitive anaerobic enzymes lasts for only a couple of days under stable conditions [36], but may be further stabilized by Fe^2+^ precipitation [37] released during the corrosion process. Because corroded infrastructure is often home to *Methanosarcina* species along with acetogenic genera, *Methanosarcina* was anticipated to play a role in Fe^0^-corrosion, however it was assumed to be indirect [7–11], so it would require a cooperation with other corrosive microorganisms, for example by retrieving the acetate produced by acetogens while corroding Fe^0^ (Fig. 1). In this study we investigate the assumption that acetoclastic methanogens like *Methanosarcina* require an interaction with an acetogen in order to corrode Fe^0^. The Bothnian Bay is an environment where corrosion could have tremendous environmental consequences. From the sediments, off the coast of Bothnia, we enriched for *Methanosarcina* on Fe^0^ [38]. A combination of scanning electron microscopy (SEM), high-throughput sequencing and physiological experiments was applied to investigate the role of methanogens and their interactions with co-occurring microbes in steel corrosion. We put forward evidence that Baltic-*Methanosarcina* can corrode Fe^0^ alone, competing with acetogens for access to Fe^0^. Based on specific inhibition experiments, we propose different mechanisms for Fe^0^ corrosion for the Baltic *Clostridium*-acetogens and the *Methanosarcina*-methanogens.

**Fig. 1.**
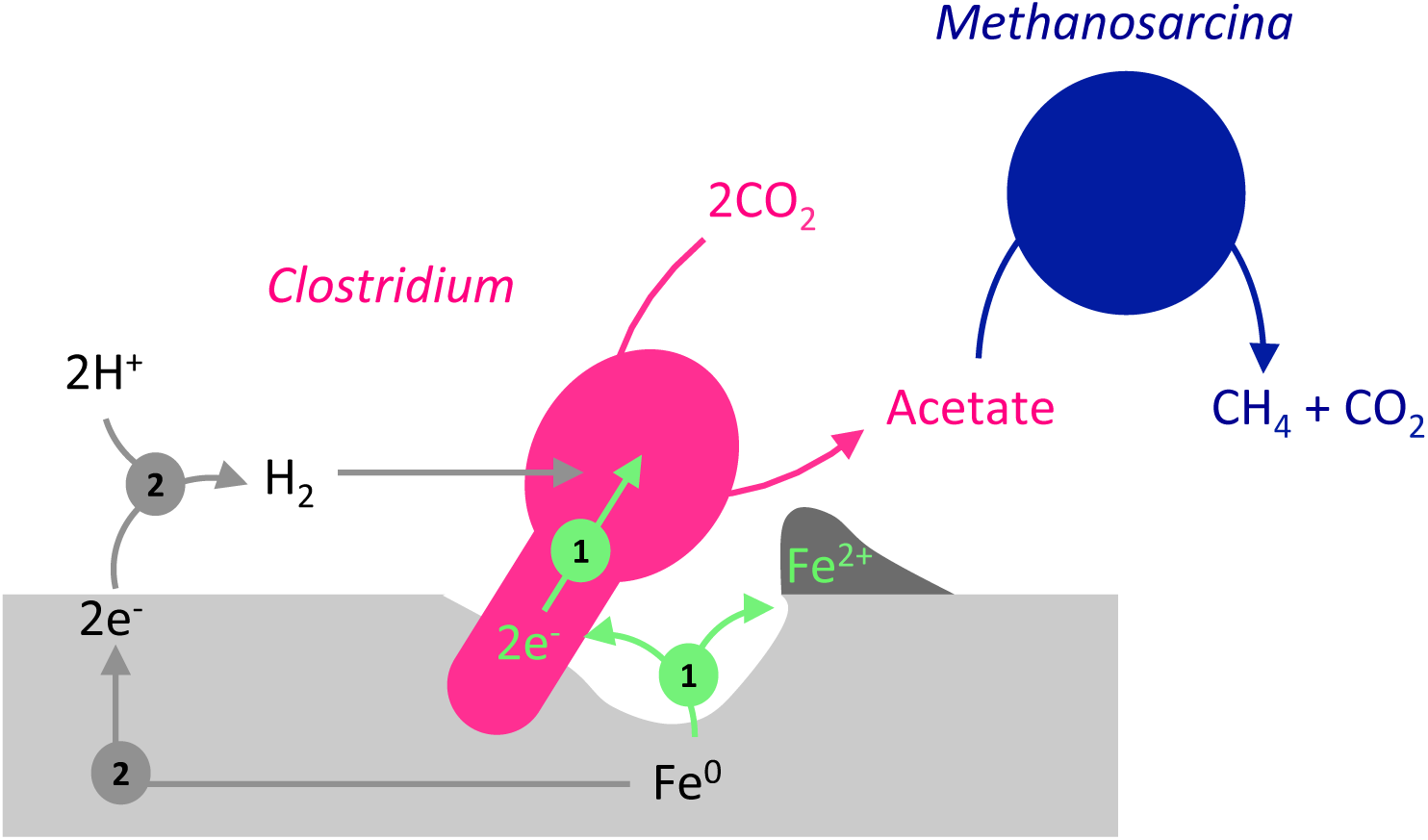
Anticipated commensal interaction between an Fe^0^-corroding *Clostridium*-acetogen and an acetate-utilizing *Methanosarcina*-methanogen. 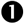Indicates a direct mechanism of electron uptake. 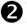Indicates a mechanism of uptake based on abiotic-H_2_.

## MATERIALS AND METHODS

### Baltic Enrichment cultures

Sediment cores were sampled during the summer of 2014 at 65°43.6’N and 22°26.8’E (station RA2) in the Bothnian Bay, Baltic Sea, at a 15 m water depth [38]. At the site the sediment had a temperature of 15°C and low in situ salinity of 0.5. The mineral content was low in insoluble manganese oxides, high in insoluble FeS, and high in crystalline iron oxides such as semiconductive goethite or conductive magnetite, as previously described [38].

Enrichment cultures were prepared using sediment from the methanogenic zone (30-36 cm) as previously described [38], but with the addition of 100 g/L iron granules, and removal of sulfide as reducing agent, which was instead replaced by an additional 2 mM cysteine (c_f_ = 3 mM). Subsequent transfers were prepared in 50 mL blue butyl-rubber-stoppered glass vials with an anoxic headspace of CO_2_/N_2_ (20:80, v/v).

Samples were enriched in a DSM120 modified medium (modifications: 0.6g/L NaCl, without casitone, sodium acetate, methanol, or Na_2_S x 9H_2_O). Iron granules (99.98%, ThermoFisher, Germany) and iron coupons (3 cm × 1 cm × 1 mm) were used as the only source of electrons and all the cultures were performed in triplicate and sometimes up to 10 replicates. Enrichments were transferred as soon as methane production reached stationary phase. Downstream analyses, DNA extractions, substrate evaluations, and SEM were performed during the fifth transfer after three years of consecutive enrichments only on Fe^0^. In addition, to confirm the presence or absence of methanogens, we used their natural, methanogen-specific autofluorescence due to coenzyme F_420_ and visualized the cells via epiflorescence microscopy.

### Chemical analyses

Methane and H_2_ concentrations were analyzed on a Trace 1300 gas chromatograph system (Thermo Scientific, Italy) coupled to a thermal conductivity detector (TCD). The injector was operated at 150°C and the detector at 200°C with 1.0 mL/min reference gas flow. The oven temperature was constant at 70°C. A TG-BOND Msieve 5A column (Thermo Scientific; 30-m length, 0.53-mm i.d., and 20-µm film thickness) was used with argon as carrier gas with a set flow at 25 mL/min. The GC was controlled and automated by the Chromeleon software (Dionex, Version 7). With this set up, the detection limit for methane and H_2_ was 5 µM.

Acetate concentrations were measured using a Dionex ICS-1500 Ion Chromatography System (ICS-1500) equipped with the AS50 autosampler, and an IonPac AS22 column coupled to a conductivity detector (31 mA). For separation of volatile fatty acids, we used 4.5 mM Na_2_CO_3_ with 1.4 mM NaHCO_3_ as eluent. The run was isothermic at 30°C with a flow rate of 1.2 mL/min. Ferrous iron in the cultures was dissolved by 0.67 M HCl (containing 0.67 M hexamethylenetetramine to avoid dissolution of metallic iron) and quantified colorimetrically using the ferrozine assay [39].

### DNA purification from microbial enrichments

DNA purification was performed using a combination of two commercially available kits: MasterPure™ Complete DNA and RNA Purification Kit (Epicenter, Madison, Wi, USA), and the Fast Prep spin MP™ kit for soil (Mobio/Quiagen, Hildesheim, Germany). 10 mL of the enrichment cultures were used for the DNA extraction, which started with the Epicenter kit with a modification to the manufacturer’s protocol: a three-fold concentration of proteinase K was added to assure cell lysis, and a prolonged incubation time at 65°C was performed until the color of the samples changed from black to brown (brown pellet gave higher DNA extraction efficiencies).

Afterwards RNase trea™ent and protein precipitation were completed with the Fast Prep spin MP^™^ kit for soil. An advantage of this kit is that it allows removal of the high iron content, simultaneously with purifying DNA on a binding matrix. DNA quality was checked on an agarose gel, and quantification took place on a mySPEC spectrophotometer (VWR^®^, Germany).

### Metagenome analyses

Metagenomic sequencing was performed via a commercially available service (Macrogen/ Europe), using an Illumina HiSeq2500 approach. Unassembled DNA sequences were merged, quality checked, and annotated using the Metagenomics Rapid Annotation (MG-RAST) server (vs. 4.03) with default parameters [40].

Illumina sequencing resulted in 10,739 high-quality reads of a total of 10,749 with an average length of 167 bp. For taxonomic analyses, the metagenomic data was compared with the Silva [41], RDP [42], Greengenes [43] and RefSeq [44] databases available in MG-RAST. The obtained rarefaction curve indicated that the prokaryotic diversity was well covered in these samples (data not shown). Deeper analyses into the microbial community structure were performed at different phylogenetic levels down to the genus level. To investigate genes involved in carbon fixation in prokaryotes and in methanogenesis, sequences were compared against the KEGG Orthology (KO) reference database. Both taxonomic and functional analyses were performed with the following cutoff parameters: e-value of 1E–5, a minimum identity of 60%, and a maximum alignment length of 15 bp. The metagenome data are available at MG-RAST with this ID: xxxxx.

### 16S rDNA sequence analyses

Archaeal primers ARC-344F (5’-ACGGGGCGCAGCAGGCGCGA-3’) [45] and ARC-1059R (5′GCCATGCACCWCCTCT-3′) [46] were used to perform PCR amplification from the isolated DNA. PCR reactions were carried in a final volume of 50 µL, which contained 1.5 mM MgCl_2_, 0.2 mM dNTPs, 0.2 µM of each primer, and 1U Taq Polymerase Promega and was completed with TE buffer. PCRs were carried out with an initial denaturation step at 94°C for 10 min; then 35 cycles of denaturation at 94°C for 30 sec, annealing at 55°C for 30 sec, extension at 72°C for 90 sec and a final extension cycle at 72°C for 10 min. 16S PCR products were cloned with the TOPO® TA Cloning® Kit for Sequencing (Invitrogen, Carlsbad, CA, USA). PCR products were sent to Macrogen Inc. (Seoul, South Korea) for Sanger sequencing using the M13F primer. Sequences were analyzed using the Geneious® software package, version 11.0.4 [47]. Sequences were compared against the NCBI GenBank DNA database using BLAST.

A consensus sequence for the Baltic-*Methanosarcina* sequences was assembled from four specific 300-600 bp sequences using ClustalW within Geneious. This consensus sequence was used to construct a maximum likelihood phylogenetic tree alongside other methanogens and a Baltic-*Methanosarcina* retrieving electrons from a Baltic-Geobacter via conductive particles [38]. Sequences were deposited in GenBank under the accession number: xxxxxx.

### Scanning electron microscopy

Iron specimens were fixed with 2.5% (v/v) glutaraldehyde in 0.1 M phosphate buffer (pH 7.3) at 4°C for 12 h, washed in phosphate buffer, dehydrated with anoxic ethanol at increasing concentrations (35%, 50%, 70%, 80%, 90%, 95%, 100%, and 3 times in 100% v/v; each step for 10 min.), then pre-dried with hexamethyldisilazane for 30 min.; [48] and dried under N_2_. Scanning electron microscopy (SEM) was performed with a FESEM Magellan 400 at 5.0 kV at the microscopy facility of the University of Massachusetts, Amherst.

### Removal of corrosion crust

Corrosion crusts from the iron coupons were removed with inactivated acid (10% hexamine in 2M HCl) [6]. N_2_ gas stream was used to dry the iron coupons, which were anaerobically stored for microscopy.

## RESULTS AND DISCUSSION

To determine if methanogenic communities from costal environments stimulate Fe^0^ corrosion we enriched for methanogenic communities from sediments offshore of the Swedish coast of the Baltic Sea. For three years we provided a Baltic methanogenic community with Fe^0^ as sole electron donor and CO_2_ as sole electron acceptor.

Original slurries (25% sediment) from Baltic sediments when provided with Fe^0^ they generated circa five times more methane and four times more acetate than incubations without Fe^0^ (Fig. 2). Methane and acetate production continued in subsequent transfers only when Fe^0^ was added as sole electron donor. After three transfers, incubations became sediment-free. In these sediment-free incubations, we noticed the formation of a black crust, which could not be observed in abiotic incubations (Fig. 2). Previously, black crust was observed under non-sulfidic conditions and classified as siderite, a common corrosion product of freshwater-microorganisms using CO_2_ as terminal electron acceptor [49]. At transfer eight, we assessed Fe^0^ corrosion, by determining ferrous (Fe^2+^) iron accumulation (Fig. 2), which is presumably generated in equimolar amounts to the Fe^0^ consumed (Fe^0^ + 2H_2_O → Fe^2+^(OH^−^)_2_ + 2H^+^ + 2e^−^), if Fe^2+^-precipitation is absent. Over the course of 25 days, the methanogenic community produced twice the amount of Fe^2+^ (1.08 ± 0.1 mM Fe^2+^) compared to abiotic controls (0.55 ± 0.08 mM Fe^2+^). Corrosion by the Baltic community was steady and significant (p < 0.0003, n>5), since corrosion started immediately and Fe^2+^ accumulated constantly above abiotic controls through the incubation. Largest difference was observed during the first five days, when Fe^2+^ was detectable only in the presence of an active community (42 ± 13 µM/day), while it was below background in abiotic incubations (<0 ± 9 µM/day).

**Fig. 2.**
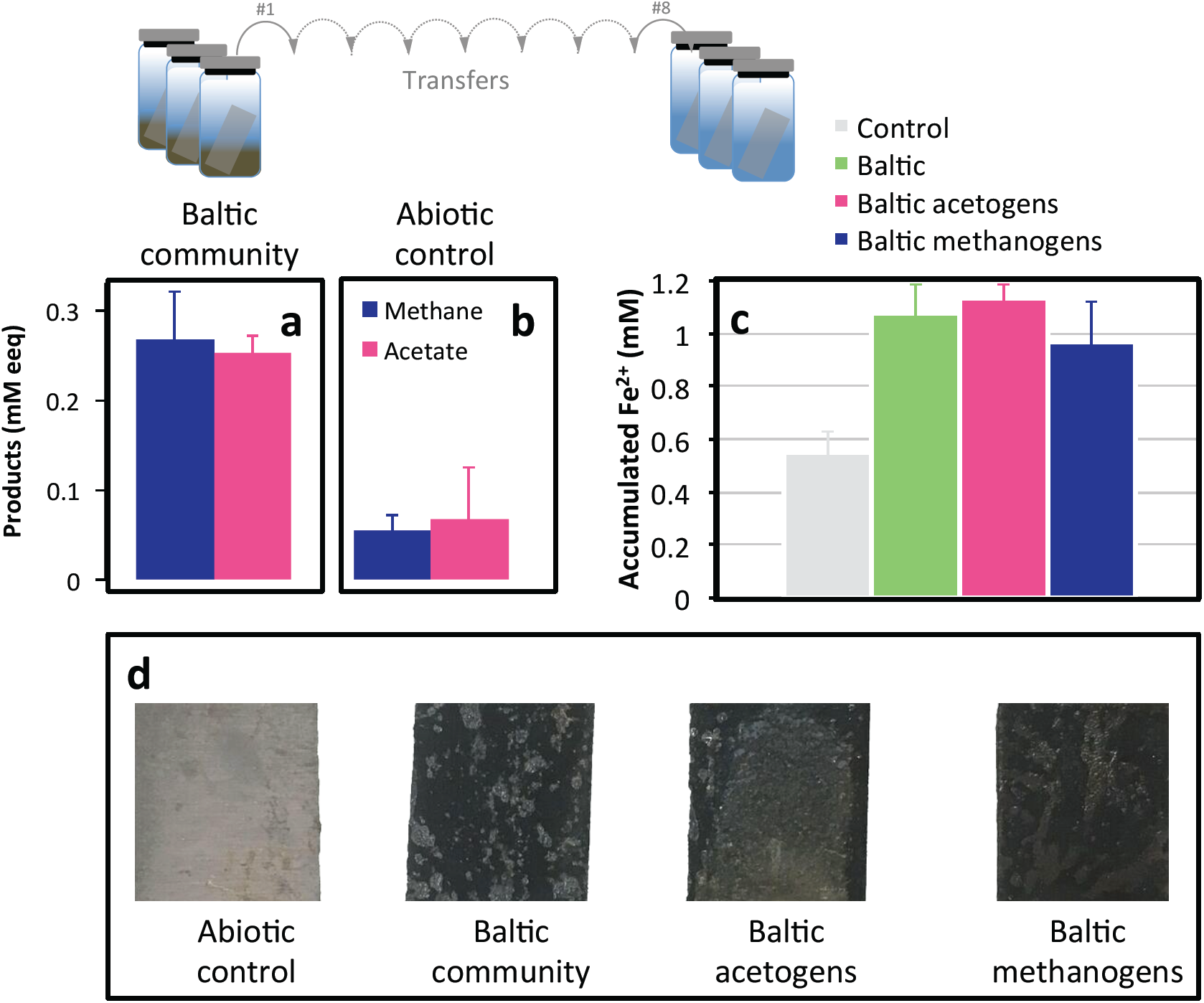
Incubations with Fe^0^ as sole electron donor in the presence (a) or absence (b) of a methanogenic community from the Baltic Sea. The top left panels show initial electron recoveries into products (methane and acetate) in the original Baltic slurries incubated with Fe^0^. Recoveries are presented as mM electron equivalents (mM eeq) considering that to make one mol methane/acetate requires 8 mols electrons (n=3). The top right panel (c) shows corrosion estimates from ferrous iron (Fe^2+^) determination. The Baltic community incubated on Fe^0^, was now at its eight consecutive transfer and accumulated Fe^2+^ to levels higher than abiotic controls, independent of the addition of inhibitors specific for acetogens and methanogens. Bottom panel (d) shows images of steel plates after three months of incubation in the presence or absence of the abiotic media, the Baltic-community, the Baltic-acetogens (by specific inhibition of the methanogens), and the Baltic-methanogens (by blocking the bacteria with antibiotics).

To determine cell types and attachment onto the metal, we carried out SEM of Fe^0^ coupons before and after three months exposure to an active Baltic community. Three major morphotypes were observed attached to the Fe^0^-coupons, a sarcina-like, a vibrio-like and rods with bulging heads, likely endospores (Fig. 3). These morphotypes resemble morphologies of *Methanosarcina, Desulfovibrio* and *Clostridium*, respectively.

**Fig. 3.**
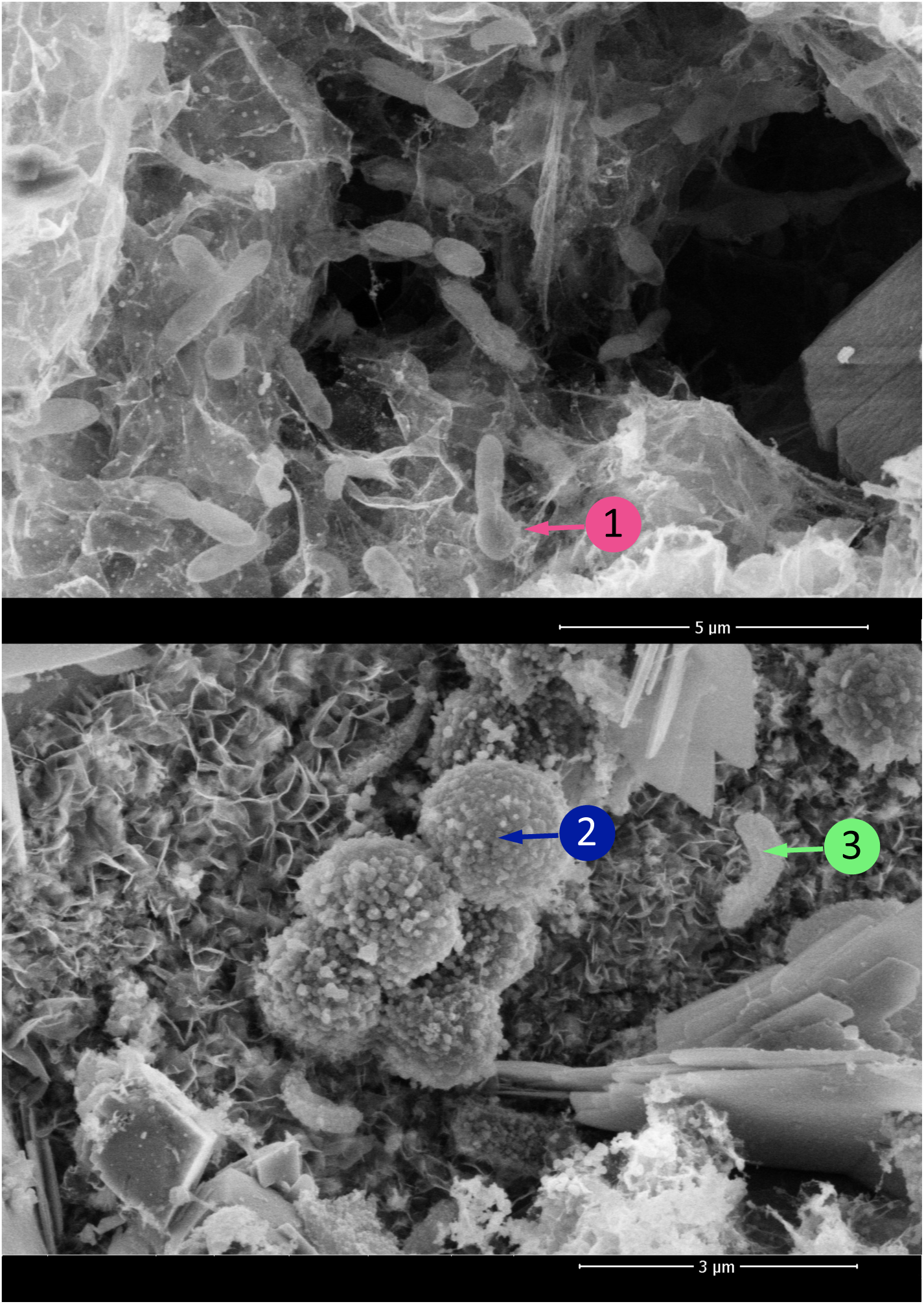
Baltic corrosive community (5^th^ transfer) established on Fe^0^-coupons for 3 months. Three morphotypes were observed: 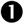 a rod-shaped cell with a bulging head – resembling a terminal endospore-forming *Clostridium*, 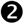 a sarcina-like cell resembling the rosette-forming *Methanosarcina* species and 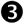 a vibrio-like cell possibly a *Desulfovibrio*.

Whole genome-based phylogenetic analyses confirmed the presence of all three groups in the corrosive enrichment (Fig. 4). There was only one dominant archaea genus, namely *Methanosarcina* (83% of all archaea; 11% of all prokaryotes). The most abundant bacterial genus was *Clostridium* (20% of all bacteria; 17% of all prokaryotes) followed by *Desulfovibrio* (6.4% of all bacteria; 5.5% of all prokaryotes). Moreover, *Methanosarcina* was the most abundant methanogen in the original sediment from the Baltic Sea, as determined by 16S-rDNA MiSeq tag-sequencing and quantitative PCR [38]. On the other hand, *Clostridium* was a minority in the Baltic methanogenic community (<1%), and *Desulfovibrio* was undetected by 16S-rDNA MiSeq of the original community [38].

**Fig. 4.**
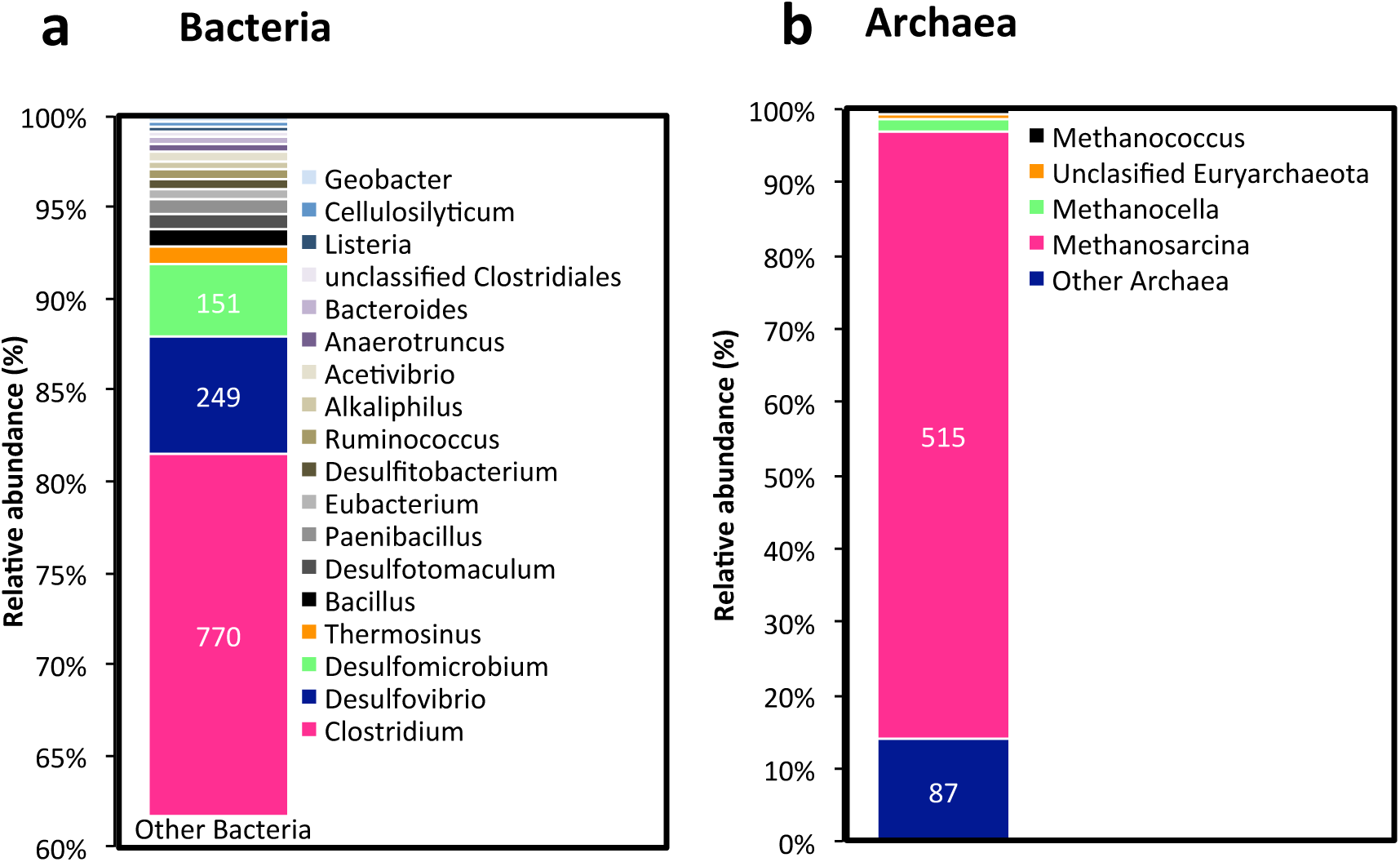
Whole-metagenome phylogeny of a community enriched on Fe^0^ for five successive transfers showing the major bacterial (a) and archaeal (b) phylotypes. For Bacteria we only show the phylotypes above 0.25% abundance. Other Bacteria and Other Archaea, include both non-abundant and unassigned species.

### Alleged commensalism between *Methanosarcina* and acetogens

The dominance of acetogenic *Clostridium*, and acetoclastic *Methanosarcina* led us to the hypothesis that *Clostridium* and *Methanosarcina* were commensals on Fe^0^ with *Clostridium* using Fe^0^ as electron donor for acetogenesis, followed by the acetate being used as food source by methanogens (Fig. 1). Thus, we expected *Clostridium* to utilize H_2_/or electrons directly from Fe^0^ and generate acetate. Few *Clostridium* species have been described to function as acetogens, like for example *Clostridium ljungdahlii* [50]. Acetogens are associated with corroded structures, and suggested to play a direct role in corrosion of Fe^0^ [7,8]. The acetate generated by acetogens is a favorable food source for *Methanosarcina* –-which would then metabolize acetate to produce methane and CO_2_ [51]. Thus, we anticipated that *Clostridium* and *Methanosarcina* live as commensals, with only the *Methanosarcina* profiting from the interaction (Fig. 1). *Methanosarcina* have been expected to play a secondary indirect role in steel corrosion [7–11]. Surprisingly, in our incubations acetate accumulated (Fig. 2), indicating that Baltic-*Methanosarcina* were ineffective acetate-utilizers. Instead they seem to use Fe^0^ as electron donor.

### Competition for Fe_0_ between methanogens and acetogens

To verify whether the Baltic *Methanosarcina* was an ineffective acetoclastic methanogen we tested its phylogenetic affiliation to a non-acetoclastic *Methanosarcina*, tested for acetate utilization, and carried out inhibition experiments in order to either block methanogenesis or acetogenesis. Corroborating our observations we determined that Baltic-*Methanosarcina*, could not utilize acetate, and therefore together with Baltic-acetogens they established a competitive-type of interaction, rather than a commensal-type of interaction.

To determine whether the methanogen was related to a non-acetoclastic *Methanosarcina* we carried out phylogenetic analyses of the 16S rRNA gene. The Baltic-*Methanosarcina* 16S rRNA-gene sequence showed 100% identity (600 bp fragment) to *M. lacustris* (Fig. 5), which unlike other *Methanosarcina*, cannot utilize acetate [52]. Once the Baltic-methanogenic community was incubated with acetate as sole electron donor, methane was untraceable for the entire incubation period, of 60 days.

**Fig. 5.**
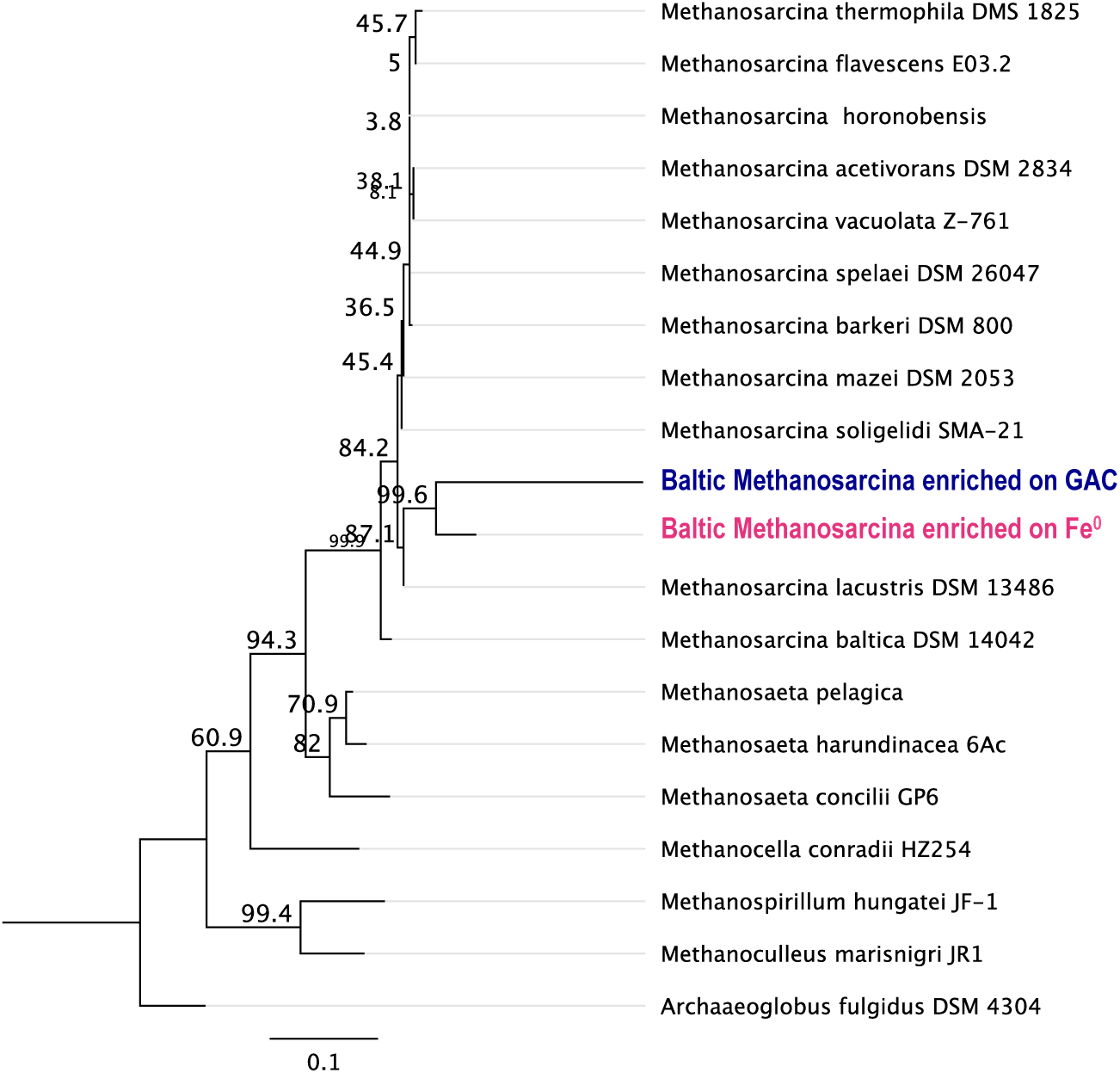
Fig. 7. Maximum likelihood tree of a 16S rRNA gene sequence of a Baltic-*Methanosarcina* capable of Fe^0^ corrosion, but incapable of acetoclastic methanogenesis over the course of 60 days. This was also aligned against a Baltic*-Methanosarcina* sequence enriched on GAC along with Baltic-*Geobacter* [38]. Scale bar represents the average number of nucleotide substitutions per site.

Inhibition of the bacterial community comprising acetogens, led to a two to six fold increase in methane-production (Fig. 6) compared to the mixed acetogen-methanogen community (Fig. 6). In the absence of acetogens (day 10-15), methanogenesis rates were higher (0.094 ± 0.018 mM/day methane) than in the mixed community (0.016 ± 0.005 mM/day methane). Antibiotics (kanamycin and ampicillin) did wear off after 15 days, resulting in a small increase in acetate production (Fig. 6), and a halving of the methanogenic rates due to the detrimental presence of ‘*undead’*-acetogens. Higher methanogenic rates when acetate-producing bacteria are incapacitated (or less active) indicate that acetogenic bacteria competitively inhibit the methanogens.

**Fig. 6.**
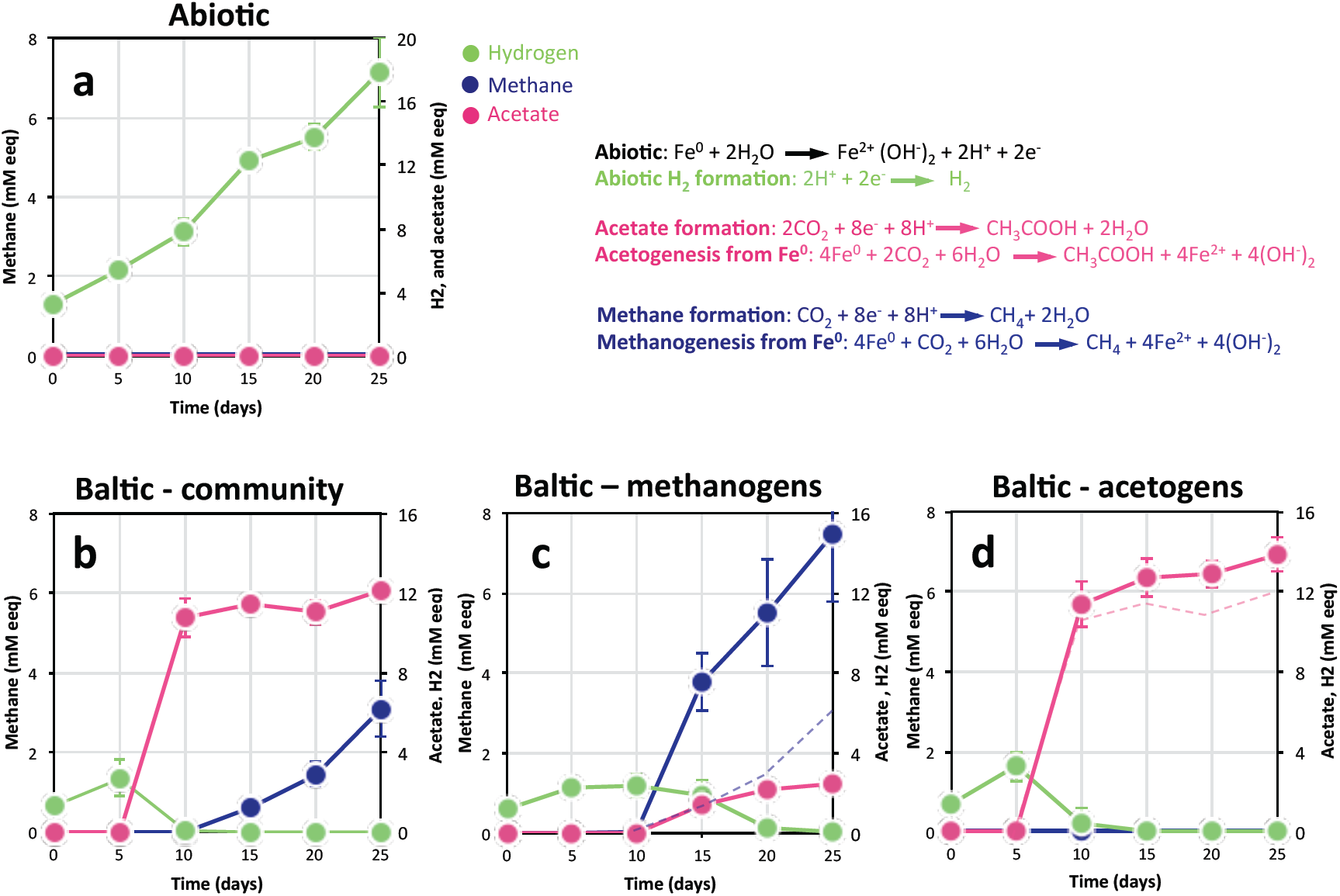
Electron recovery (as mM electron equivalents) in products, considering that 2 mM electrons are required to make 1 mM H_2_ and 8 mM electrons are required to make 1 mM acetate or methane according to the reactions in the right upper panel. (a) In abiotic controls electrons are recovered as H_2_. (b) The whole community grown on Fe^0^ for eight transfers recovered electrons as methane and acetate. (c) Inhibition of all bacteria (including acetogens) with a cocktail of antibiotics (kanamycin and ampicillin) led to the survival of methanogens, which produced more methane than the mixed-Baltic community (dotted line/b) (d) Specific inhibition of methanogens with 2-bromoethanesulfonate, led to the persistence of a community of acetogens, which produced more acetate than the mixed-Baltic community (dotted line/b). (n>5)

Furthermore, to determine whether acetogens were negatively impacted by methanogens we inhibited the methanogenic community with 2-bromoethanesulfonate (BES), a methyl–CoA analogue. Using BES as inhibitor, the methanogenic community was rendered inactive for the entire period of the incubation (Fig. 6). Acetogens alone were significantly more productive (14%; p<0.0001) than acetogens in the mixed community. Therefore we could conclude that within the mixed community methanogens did pilfer access to Fe^0^ from acetogens in their struggle to survive (Fig. 6). Thus both, acetogens and methanogens were negatively affecting one another when competing for Fe^0^ as sole electron donor (Fig. 7).

**Fig. 7.**
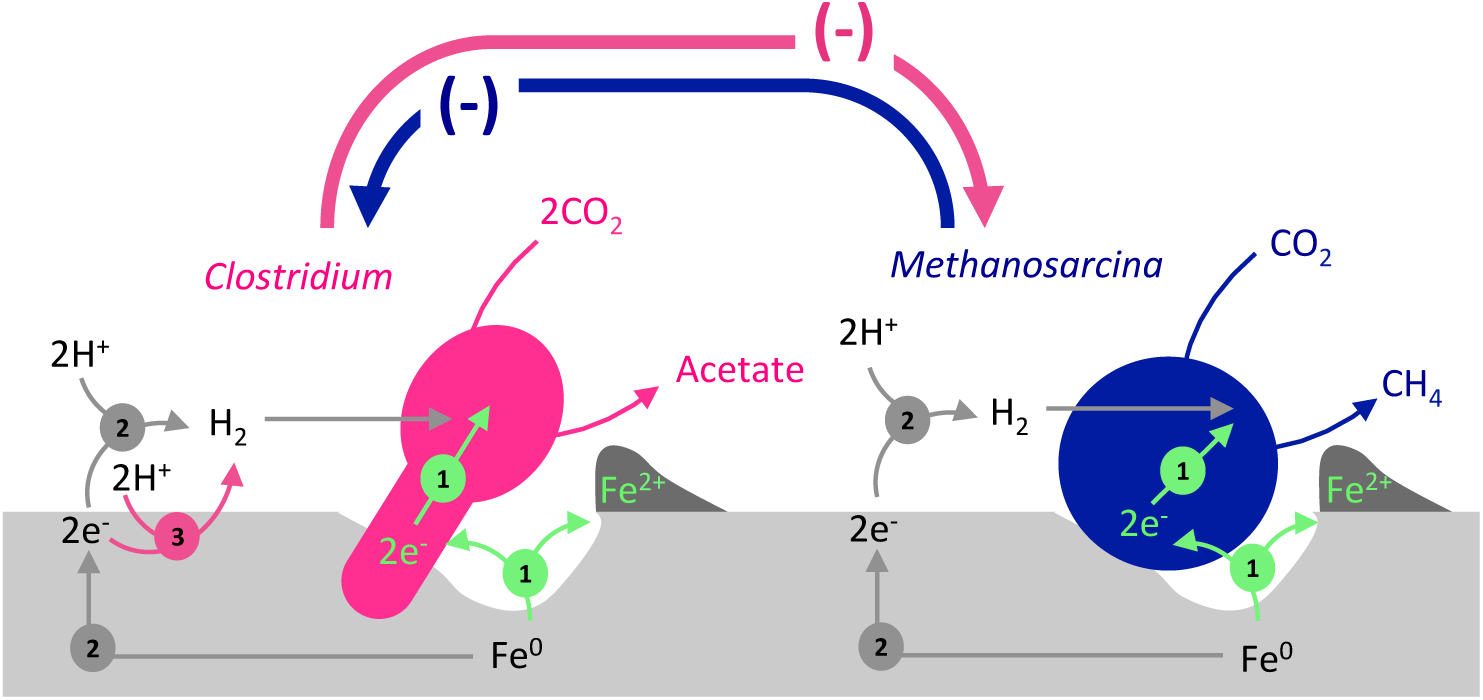
Model of the competitive interaction between an Fe_0_-corroding *Clostridium*-acetogen and an Fe_0_-corroding *Methanosarcina*-methanogen. 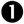Indicates a direct mechanism of electron uptake. 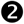Indicates a mechanism of uptake based on abiotic-H_2_. 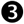Indicates extracellular enzyme-mediated H_2_-evolution.

Here, for the first time, we bring evidence that an environmental *Methanosarcina* is directly involved in Fe^0^ corrosion. Plus our results show that methanogenic activity in a *Methanosarcina* dominant Baltic community does not depend on the bacteria or the substrates they generate. Our results are contesting previous suppositions that *Methanosarcina* and acetogens mainly interact syntrophically, via acetate-transfer, within a corrosive community [53].

### Mechanisms of electron uptake from Fe^0^

Acetogens have been credited to use a variety of mechanisms for electron uptake from Fe^0^ including uptake of abiotic H_2_ [54], uptake of enzymatically evolved H_2_ [35,55] or direct-electron uptake. The later is a possibility inferred from the property of some acetogens to grow on electrodes under non-hydrogenogenotrophic conditions [50, 56, 57].

Here we demonstrate, that Baltic-acetogens are most likely using oozed endogenous enzymes for quick retrieval of electrons from Fe^0^. This was similar to earlier studies, which investigated pure culture acetogens corroding Fe^0^, such as *Sporomusa* or *Acetobacterium* strains [35,54].

In our study, acetogenesis started 5 days earlier when we added a filtrate of spent-media from a Fe^0^-grown Baltic community (Fig. 8). Plus, acetate recoveries were higher at the addition of the spent filtrate (Fig. 8). We observed that accumulation of acetate increased by 22% (n=10, p<0.00001) compared to the mixed Baltic community, and by 7% (n=10, p<0.02) compared to Baltic-acetogens alone, after inhibiting their competitors, the methanogens. We conclude that Baltic-acetogens are most likely to use a mechanism of electron uptake from Fe^0^ mediated by enzymes similar to other acetogens [35,54]. In the Baltic corrosive community *Clostridium*-species are likely to play the role of acetogens, since several species have been shown to produce acetate either by electrosynthesis of autotrophically on H_2_ [50, 58, 59]. Besides, clostridial enzymes, for example [Fe]-hydrogenases from *Clostridium pasteurianium,* were shown capable of corrosion on their own, assumedly by direct retrieval of electrons coupled to proton reduction to H_2_ [60,61]. *If* a sacrificial population exuded enzymes, *then* the addition of spent-media filtrate, expected to contain exuded enzymes, would significantly stimulate H_2_-production and subsequent H_2_-uptake by Baltic-acetogens. This was not the case, as indicated by comparable H_2_-concentrations in incubations with Baltic-acetogens with or without additional endogeneous enzymes from spent-filtrate (1.3 ± 0.5 mM versus 1.6 ± 0.4 mM respectively; n=10; p=0.4).

**Fig. 8.**
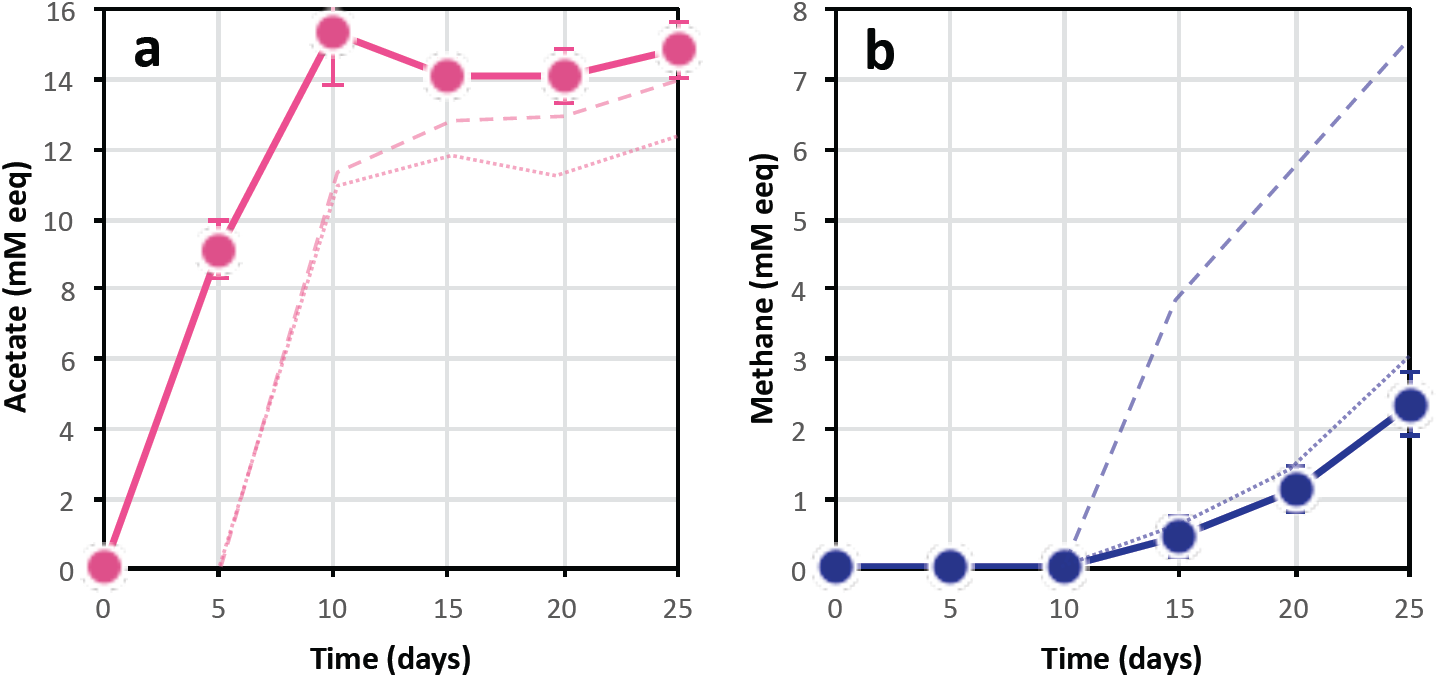
Impact of spent-filtrate spiking on acetogenesis (a) and methanogenesis (b) in freshly-prepared Fe^0^-Baltic communities (n=10). Dashed lines show the trends for product formation by the acetogens-alone and methanogens-alone when their competitors were specifically inhibited. Dotted lines show the trends for product formation when the acetogens and methnaogens were competing with one another.

On the other hand, methanogens showed a 23% drop in methane productivity (n=10, p<0.03; Fig. 8) at the addition of Fe^0^ spent media filtrate as compared to the mixed Baltic community. It is therefore unlikely that *Methanosarcina* is using an enzymatic-mediated electron uptake mechanism. These data were also corroborated with a previous study on Baltic-*Methanosarcina* capable of mineral-syntrophy independent of enzyme, as it remained unaffected by spent filtrate additions [38].

Although it has been suggested that enzyme-mediated corrosion may occur in methanogens, this property has only been demonstrated for *Methanococcus*-species [35,55]. Initial studies looking at *Methanosarcina*’s ability to corrode Fe^0^ assumed these methanogens were using abiotic H_2_. Nevertheless, *Methanosarcina* have high H_2_-uptake thresholds (296 nM - 376 nM) [62,63], and *Methanosarcina* should theoretically be outcompeted by other methanogens with lower H_2_-uptake thresholds (e.g. circa 6 nM for *Methanobacterium formicicum* [62,63]. This is not the case. *Methanosarcina* are often found associated with Fe^0^ corrosion rather than hydrogenotrophic methanogens with lower H_2_-tresholds. Plus in our system, Baltic-*Methanosarcina* did not benefit from spent-filtrate addition, which is expected to include hydrogenase enzymes [35,54]. Instead, Baltic-*Methanosarcina* may be directly reclaiming electrons from Fe^0^ as they do from other cells [25,26] or conductive particles [25, 29, 38], consequently stealing electrons from competing H_2_-utilizers, methanogens or acetogens, within a corrosive community. The mechanism of direct electron uptake in *Methanosarcina* has been only recently challenged using a comparative transcriptomics approach of *Methansarcina* provided with electrons directly from a current-producing syntrophic partner (*Geobacter* [25,26]) or with H_2_ from a H_2_-producing syntrophic partner (*Pelobacter*). For a direct-type of interaction, it was previously shown that *Methanosarcina* up-regulates cell-surface proteins with redox properties such as cupredoxins, cytochromes and other Fe-S-proteins [34].

However, the exact role of these redox-active proteins in how *Methanosarcina* retrieves electrons from solid extracellular electrons donors (Fe^0^, other cells or, electrically conductive particles) remains enigmatic and deserves future exploration.

## CONCLUSION

*Methanosarcina* and acetogens are often found on corroded Fe^0^-structures in non-sulfidic environments. However, the role of *Methanosarcina* was assumed to be rather secondary, as a commensal feeding on the acetate produced by acetogens implicated directly in corrosion. Here we investigate a corrosive methanogenic community from the Baltic Sea, where steel corrosion of chemical weapons, radionuclide waste containers and pipeline structures are an economic and environmental threat.

*Clostridium*-acetogens and *Methanosarcina*-methanogens dominated the corrosive Baltic Sea community. Our results demonstrated that Baltic-*Methanosarcina* does not establish a syntrophic interaction with the acetogens as often reported. Instead, *Methanosarcina* and the acetogens compete with each other to reclaim electrons from Fe^0^. While Baltic acetogens seem to use endogenous enzymes, *Methanosarcina* were not. We suggest Baltic-*Methanosarcina* may be retrieving electrons directly via a mechanism that is largely unexplored.

## AUTHOR CONTRIBUTIONS

PAP and AER designed the experiments. AER carried out the sampling, and processing of the Baltic sediment as well as the original incubations with help from BT. PAP carried all downstream growth experiments and analyses. PAP did all molecular experiments and analyses with support from OSW and CL. PAP and AER wrote the manuscript and all authors contributed to the final version of the manuscript.

## ACKNOWLEDGMENTS

This is a contribution to a Sapere Aude Danish Research Council grant number 4181-00203 awarded to AER. We would like to thank Lasse Ørum Smidt, Erik Laursen, and Heidi Grøn Jensen who were instrumental with lab assistance through the project.

We thank for assistance Joy Ward and the microscopy facility unit at the University of Massachusetts Amherst who assisted us with scanning electron microscopy.

